# Mild impacts do not subsidize aquatic insect communities in an Atlantic Forest stream

**DOI:** 10.1101/2021.01.24.428003

**Authors:** Celio Roberto Jonck, Luiza Hoehne Mattos de Oliveira, Rafael Jordão Pires Silva, Jorge Luiz Nessimian

**Affiliations:** Leopoldo Americo Miguez de Melo Research Center, Rio de Janeiro, RJ, Brazil; Zoology Department, Federal University of Rio de Janeiro – UFRJ, Rio de Janeiro, RJ, Brazil

**Keywords:** aquatic insects, organic pollution, toxic pollution, subsidy-stress gradient

## Abstract

Odum’s perturbation theory hypothesizes that toxic pollutants cause damage to ecosystems early in the course of contamination. In contrast, organic pollutants enrich the ecosystem until it exceeds their carrying capacity, an effect known as the subsidy-stress gradient. Understanding this dynamic can improve the efficiency of river restoration programs and bring significant benefits to society by providing ecosystem services that were lost. However, the initial effects of the most common human-induced disturbances in Atlantic Forest streams are not well known, indicating the necessity to evaluate the subsidy-stress gradient in these vulnerable and diverse ecosystems.

**Aim:** We evaluated the composition and abundance of the community of aquatic insects from leaf litter of headwater streams in three conditions: a fully forested area (reference stream), a low-intensity urban settlement (urban stream), and a region with small farms dedicated to the cultivation of fruits and vegetables (agricultural stream).

**Methods:** We used alpha and beta diversity metrics and a specific biotic index to test the subsidy-stress gradient prediction.

**Results:** The agricultural stream showed the most degraded ecological condition. The urban stream and the reference stream showed similarity in alpha diversity metrics. According to the biotic index, the streams showed a gradient of environmental quality, with the reference stream showing the best quality and the agricultural stream the worst quality.

**Conclusions:** The agricultural stream showed a decrease in the environmental quality consistent with the effect predicted by the subsidy-stress gradient due to toxic pollutants’ contribution. However, the low-intensity enrichment of organic matter from the urban settlement causes a disorder in the ecosystem that reduces its environmental quality, contrary to the predicted by the subsidy-stress gradient.

## 1. Introduction

Land use has become an essential issue for environmental conservancy since meeting the human needs for food, shelter, and energy requires the intensive use of large areas and ends up threatening the very capacity of ecosystems to provide environmental services (Foley et al., 2005).

Brazilian population intensively uses the Atlantic Forest as a natural resource provider (Dean, 1996), which decimated its original coverage and left remnants highly fragmented and poorly protected (Ribeiro et al., 2009).

Therefore, two of the most common impacts in this biome are urban settlements and agricultural activities. Urban settlements use less land but may cause more impact depending on the city’s size and its capacity to perform sewage treatment (Fistarol et al., 2015, Fernandez-Cassi et al., 2018). In contrast, agricultural activities are punctually less intense but require a more substantial area that tends to grow when production is inefficient (Vandermeer & Perfecto, 2007). In general, this forms a gradient in which preserved environments have higher environmental quality than agricultural areas, which have higher environmental quality than urban areas (Stranko et al., 2012).

Due to impacts on the environment triggered by human society, several authors predict problems for our economy (Tol, 2018), our mental health (Evans, 2019), and even our survival (Bonan & Doney, 2018). Meanwhile, environmental monitoring continues to have to convince the world that it is necessary (Lovett et al., 2007), postponing the development of a discipline with predictive capabilities, less based on damage documentation, and more focused on prevention (Cairns Jr. 1981).

Understanding how each disturbance affects ecosystems is essential for developing a predictive, robust, and well-defined impact mitigating science. One of the first proposals on this topic was the Perturbation Theory and the Subsidy-Stress Gradient (Odum et al., 1979). This theory predicts that when a pollutant is toxic to the ecosystem, species richness tends to fall early in the course of contamination. On the other hand, when the pollutant is usable as a resource by biological communities, it will increase ecosystem performance (Odum et al., 1979) and enrich the environment until it reaches its carrying capacity, which will lead to a population decline.

Odum et al. (1979) conceive the subsidy-stress gradient from a macro-scale perspective since the examples they use for the phenomenon refer to ecosystem responses to disturbances (e.g., river swamps’ response to flood duration). However, later the primary author uses the term stress to denote a detrimental influence and the term subsidy for a positive influence, although followed by negative responses (Odum, 1985). This statement suggests that the subsidy effect could occur on a microscale, although the disturbance has adverse effects on the ecosystem.

Besides, the classification of the response as positive or negative is also subject to interpretation. A good example is an increase in community respiration. Odum himself classified that as the first early warning sign of stress (Odum, 1985), while other authors classified that as a pure subsidy effect (Pereda et al., 2019). Precise definitions of scale and classification of responses are needed to avoid misunderstandings on applying the subsidy-stress gradient knowledge to ecosystems conservation or restoration.

Whatever the degradation processes, they tend to strongly affect the lotic environments since the valley has a significant influence on the characteristics of the stream (Hynes, 1975; Sioli, 1975), and even changes that occur in the terrestrial environment or the atmosphere can affect them (Williamson et al., 2008). Nowadays, we understand rivers as four-dimensional systems interconnected with the whole landscape. They integrate the conditions of drainage from headwaters to mouth, the input of organic matter from the adjoined land, the interaction of surface waters with groundwater, and changes over time superimposed on all the spatial dimensions (Ward, 1989). While integrative capacity may be a problem for a river’s environmental quality, it is also the main reason why they are always considered promising targets for environmental monitoring (Karr, 1998).

Headwaters show high habitat heterogeneity, produced mainly by the dynamics of transport and sedimentation of mineral substrates and plant material deposition. This heterogeneity is essential for structuring benthic macroinvertebrate communities (Minshall & Robinson, 1998). By joining the functions of shelter and especially food (Richardson, 1992), the leaf litter ends up being colonized by a large number of organisms that form a reasonably structured community (Wallace et al., 1997), which is why many studies use them for water quality assessment (Medved, 2013).

Here we test whether the effect hypothesized by the subsidy-stress gradient occurs in the community of leaf litter aquatic insects of an Atlantic Forest stream subjected to mild impacts of both toxic and usable pollutants. We are seeking to understand the response of these environments to early signs of human occupation.

## 2. Methods

### 2.1. Study Sites

We perform the study in the Guapiaçu River Basin headwaters, located at the Serra do Mar mountain range between Teresópolis and Cachoeiras de Macacu, Rio de Janeiro. Based on the watershed’s land use, three sampling points were defined. The first sampling point, located at 22°25′05.2″S and 42°44′19.2″W, had no visible human occupation. This point received no input from contaminants, and we defined it as the reference stream. The second sampling point, located at 22°29′25.5″S 42°50′02.3″W, had part of the valley occupied by small farms, which is why we define it as the agricultural stream. Although the cultivation area was small, we confirmed that farmers used pesticides such as Roundup, Kumulus, and Phosmet. Therefore, it is plausible to assume that this point was under the influence of toxic input. The third sampling point, located at 22°26′51.6″S 42°47′17.6″W, had part of the valley occupied by housing, small businesses, and few areas with subsistence farming, which is why we define it as the urban stream. Due to the upstream human presence, it is plausible to assume that this point received organic (usable) input.

### 2.2. Field and Laboratory Procedures

We collected five samples of the substrate formed by organic matter accumulations (leaf litter). For this, we used a Surber with 30 cm² of capture area and a mesh with a 0.25 mm aperture. The collected material was washed in running water, sorted, and identified at the family level under stereomicroscope and identification keys of aquatic insects from Rio de Janeiro lotic environments (Mugnai et al., 2010).

### 2.3. Data analysis

The specimens were counted and separated by families since this was the taxonomic level required by the indexes we use. All subsequent calculations were based on this taxonomic level and executed in the R environment (R Core Team, 2019). We use the vegan package (Oksanen et al., 2019) to measure Shannon, Pielou, and Fisher’s alpha indices (Fisher et al., 1943) and the sample-based rarefaction (Gotelli & Colwell, 2001). We use the SPECIES package (Wang, 2011) to estimate Jackknife (Burnham & Overton, 1978) and ACE (Chao & Lee, 1992) total richness, the ineq package (Zeileis & Kleiber, 2014) to measure the Gini index (Cowell, 2000) and the rareNMtests package (Cayuela & Gotelli, 2014) to measure individual-based rarefaction (Gotelli & Colwell, 2001). We use packages ggplot2 (Wickham, 2016), dendextend (Galili, 2015), and ggrepel (Slowikowski, 2018) to construct graphics and Google Maps (Google, USA) to obtain satellite images. Raw data and calculation codes are available in SM1.

We also calculated the Leaf Pack Network (LPN) biotic index, an initiative that uses the colonization of leaf packets to monitor streams’ quality (Medved, 2013). As we only evaluated aquatic insects, other groups were considered absent for the calculation of the index. Therefore, the values presented here should be used only to compare the sampling stations of this study, not been comparable with the degree of organic pollution of LPN quality reports or other studies carried out with this method. We show LPN quality reports in SM2. Finally, we analyzed the beta diversity with a non-metric multidimensional scaling and a cluster analysis, both using the dissimilarity of Bray-Curtis in the scaled data.

## 3. Results

A total of 2248 individuals belonging to 37 families were collected. The most abundant families were Chironomidae, Leptoceridae, Simuliidae, Elmidae, and Philopotamidae (Figure 1).

**Figure 1.**
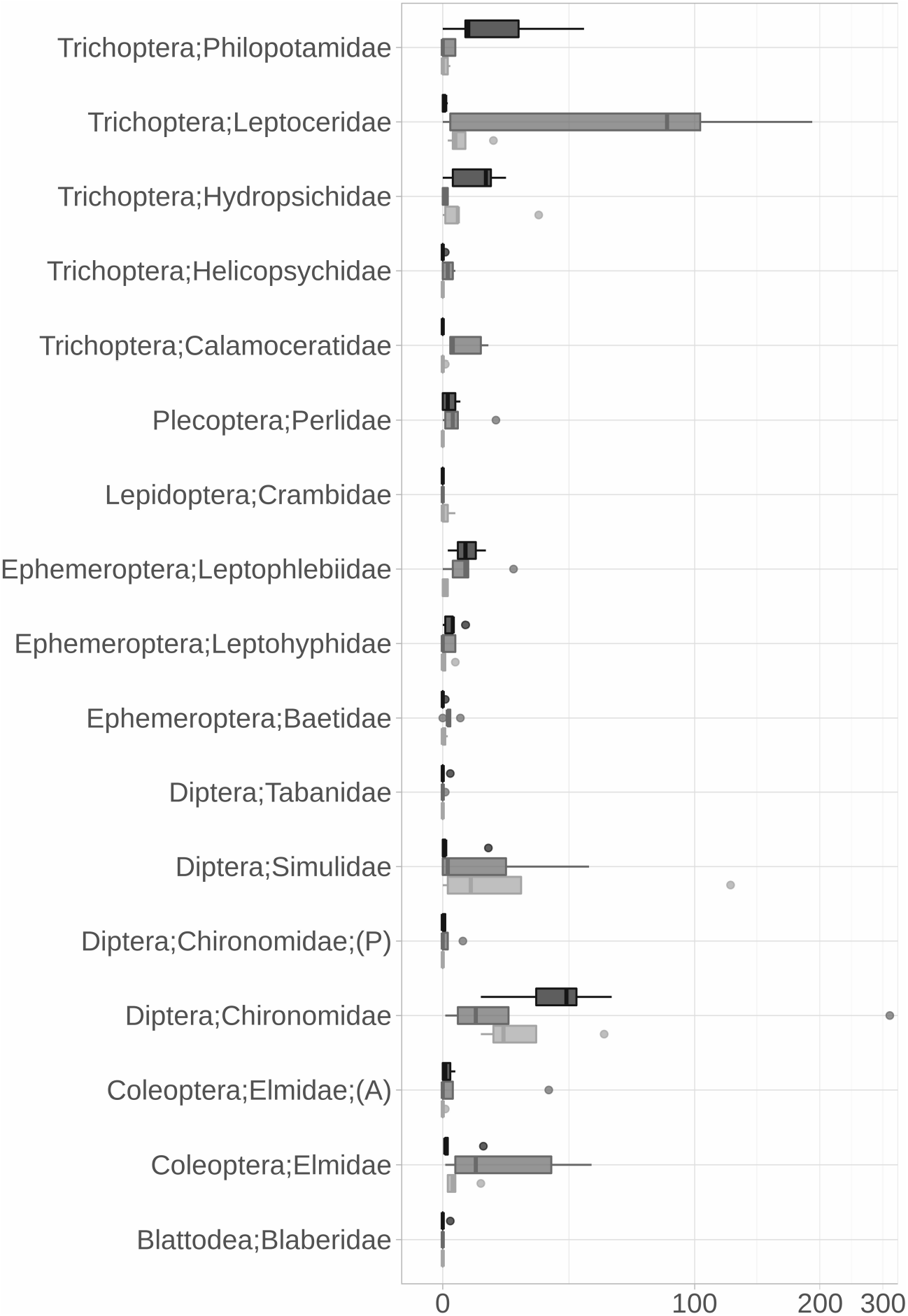
Family composition by number of individuals. Only families with abundance greater than three are represented. Dark gray is urban stream, light gray is reference stream, and medium gray is agricultural stream. Black bar in the middle of the boxplot is the median. A – adult, P – pupae.

Leptoceridae and Elmidae were more abundant in the reference stream than in disturbance streams. Philopotamidae dominated the urban one and Simuliidae the agricultural one. Chironomidae had the lowest median abundance in the reference stream, but one replica had more than 300 individuals. In disturbance streams, the distribution of this family was more balanced.

We present the absolute richness values and the total richness estimators in table 1. The reference stream showed a higher number of individuals, richness, and total richness prediction by the ACE method. The urban stream was similar to the reference in all metrics except for the number of individuals. The agricultural stream had a smaller number of individuals but still had a higher predictive value of total richness by the Jackknife method.

**Table 1.** Absolute and estimated total richness based on family-level taxonomy.

|  | Reference | Agricultural | Urban |
| --- | --- | --- | --- |
| Individuals | 1215 | 486 | 549 |
| Richness | 26 | 18 | 25 |
| Singles | 7 | 8 | 7 |
| Doubles | 3 | 0 | 6 |
| Jackknife | 33 | 34 | 32 |
| ACE | 35 | 30 | 34 |

Using the rarefaction method, we calculated the normalized richness by the number of individuals (480) for each stream. In this case, the urban stream was the richest with 24 families, followed by the reference stream with 20 families and the agricultural stream with 18 families (Figure 2A). This result is severely affected by the large number of individuals from a single-family and by the normalization process in the reference stream, as can be seen by the considerable distance between the normalization line and the end of the dark gray curve in figure 2A (reference stream). As the number of samples was identical in all the streams sampled, the sample-based rarefaction curve is not affected by this normalization, nor is it influenced by the number of individuals. In this case, the reference stream was the richest with 26 families, followed by the urban stream with 25 and the agricultural stream with 18 families. We show sample-based rarefaction curves in Figure 2B.

**Figure 2.**
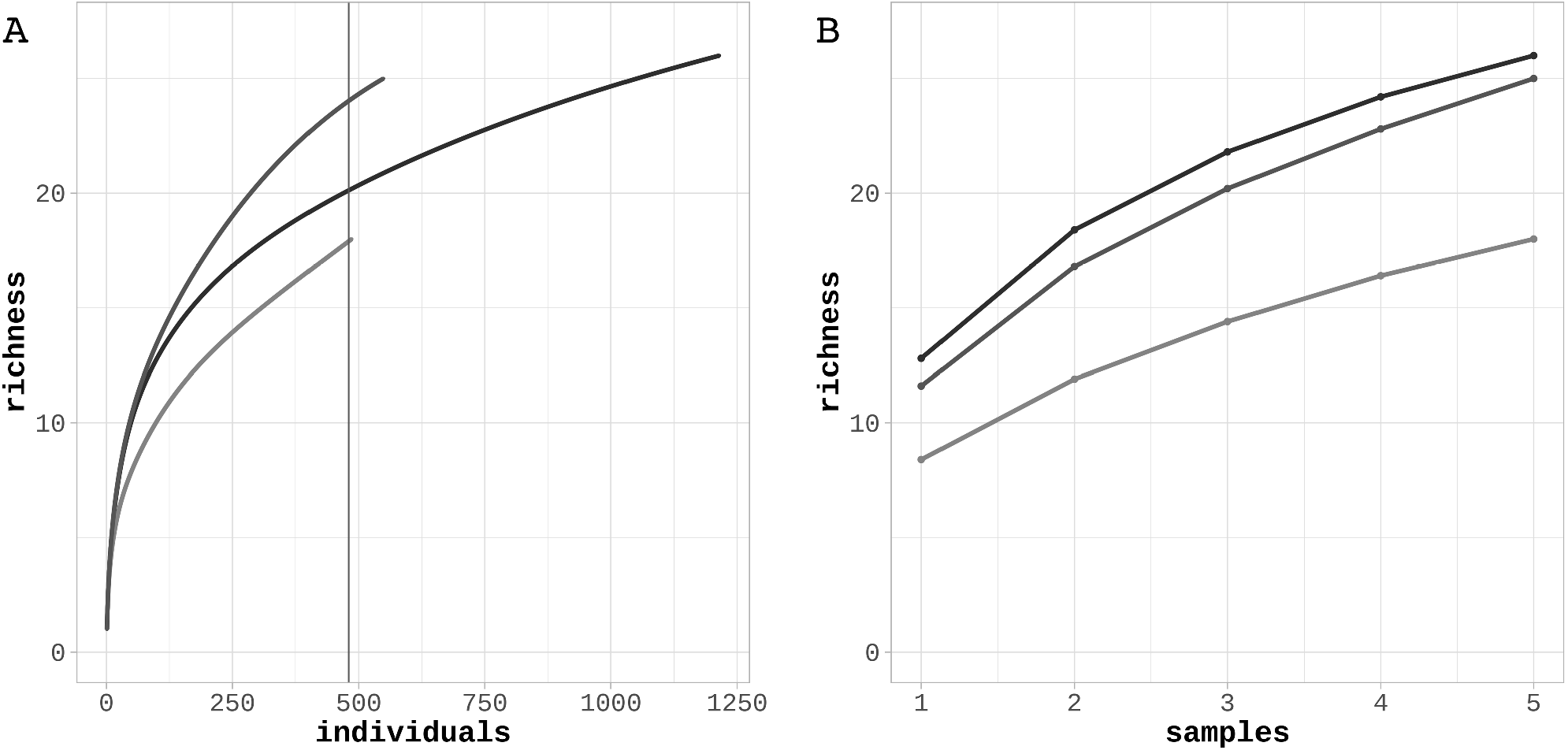
Rarefaction (A) and accumulation (B) curves. The vertical line in rarefaction curve is the subsample we use to normalize richness (480). Dark gray line is reference stream, light gray line is agricultural stream and medium gray line is urban stream.

The Shannon diversity index showed that urban and reference streams are very similar and with greater diversity than the agricultural stream, the same pattern presented by %EPT. The Fisher’s Alpha showed a gradient with the urban stream showing greater diversity, followed by the reference stream and lastly the agricultural stream. The three streams showed similar evenness and inequality.

In contrast, the LPN biotic index showed a gradation with the reference stream with the best conservation status and the agricultural stream with the worst, with the urban stream in an intermediate position. All alpha diversity and biotic indexes are shown in table 2.

**Table 2.**
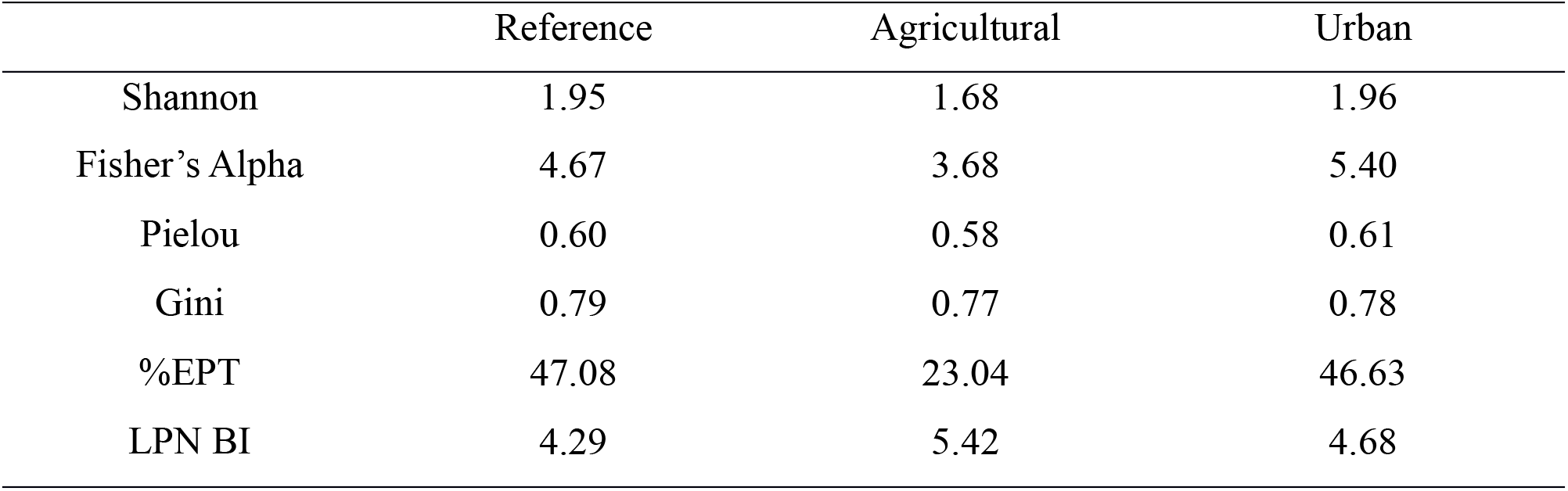
Alpha Diversity and Biotic indices

In the beta diversity cluster analysis, we verified three groups’ formation with a tendency for micro watershed separation. However, a sample of the urban stream grouped with the agricultural stream and two samples of the reference stream formed a subgroup closest to the urban stream samples. There was a significant dissimilarity between replicates, with the reference stream having the most similar replicates. Even so, the similarity between the closest replicates did not reach 50% (Figure 3A). We observe a more precise representation in the multidimensional scaling. Complete separation of the samples from the reference stream and the agricultural stream is evident, with the samples from the urban stream interspersed between them (Figure 3B).

**Figure 3.**
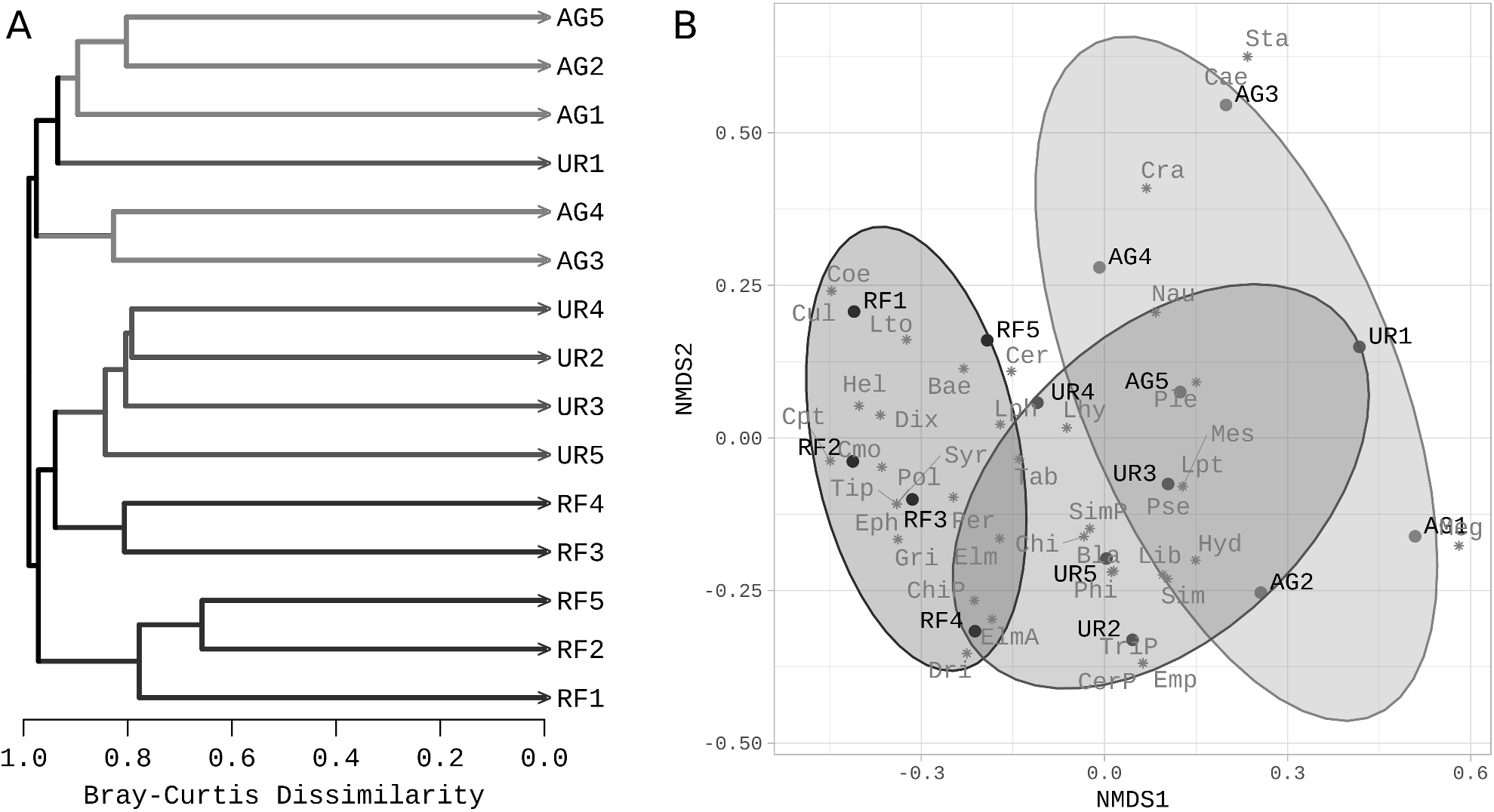
Beta diversity analysis based on Bray-Curtis dissimilarity. (A) Cluster analysis and (B) Non-Metric multidimensional scaling. AG: agricultural stream; UR: urban stream; RF: reference stream; The number after the code is the replicate number.

## 4. Discussion

The reference stream presented more individuals than the combined urban and agricultural streams (Table 1), and Leptoceridae was the most notable family, making up almost one-third of the sample (Figure 1). This dominance could raise the suspicion that the reference area only has more individuals because it has more dominance than the other areas. However, the Pielou’s evenness and Gini’s inequality indices did not show considerable differences between the communities (Table 2), indicating that dominance is a common feature of the streams. The reference stream only has a higher amplitude of dominance because it has more individuals than the other streams. The Philopotamidae and Simulidae families dominated the urban and agricultural streams, respectively (Figure 1).

Applying the BMWP score adapted to Brazilian streams (Loyola, 2000) to the dominant groups, we can see that the reference stream is dominated by a family of Trichoptera (Leptoceridae) with the highest sensitivity score (10 out of 10). A family of Trichoptera also dominated the urban stream, but in this case, it was Philopotamidae, which is more resistant to pollution (8 out of 10) than Leptoceridae. In turn, the agricultural stream presented Diptera (Simuliidae) as dominant. The entire Diptera order is considered more resistant to pollution than Trichoptera, but within the group, Simulidae is one of the most sensitive families (5 out of 10), which indicates that impacts are not too severe.

Besides the dominant ones, we can see that the composition of the reference area has a more significant number of families with high sensitivity to pollution. In addition to Leptoceridae, we found the families Calamoceratidae (10 out of 10) and Elmidae (5 out of 10, but the most sensitive family within Coleoptera) with considerable abundance. In urban and agricultural streams, after Philopotamidae and Simuliidae, the most abundant groups were Hydropsichidae (5 out of 10) and Chironomidae (2 out of 10).

The Chironomidae family presented a rather unusual distribution in the reference stream. A replicate completed more than 300 specimens, while the median was less than 15, which probably reflects an aggregate distribution pattern. However, both streams with disturbances did not show the same pattern since this family’s distribution was more balanced but had higher medians (Figure 1). Density variations in aquatic insect populations are not uncommon (Downes et al., 1993). The presence of a high-density replica of Chironomidae should be analyzed sparingly concerning the level of pollution tolerance attributed to the group (Marques et al., 1999) since the capacity to withstand a higher concentration of pollutants cannot be confused with the inability to live in clean environments.

The aspect of the community that most approached the response curves hypothesized by the subsidy-stress gradient was diversity. Absolute richness values, total richness estimators (Table 1), sample-based rarefaction curve (Figure 2B), Shannon diversity index and EPT percentage (Table 2) showed the urban stream considerably similar to the reference stream. The individual-based rarefaction curve (Figure 2A) and Fisher’s Alpha diversity index (Table 2), in turn, responded as hypothesized by the subsidy-stress gradient, indicating that the urban stream would be more diverse than the reference stream. The agricultural stream had the lowest values in all cases.

Although diversity is widely used in ecological studies (Magurran, 1988) and pointed as an indicator of ecosystem stability (Loreau & de Mazancourt, 2013), it can be confused by intrinsic environmental characteristics, such as the higher amplitude of dominance verified in the reference stream. This dominance is the likely cause of the difference presented by the individual-based and sample-based curves. The LPN biotic index was constructed to evaluate differences in stream quality based on the macroinvertebrate community composition that colonizes the leaf litter of headwater streams. Therefore, this was the tool most closely dedicated to answering the central question of this study. Its biotic index is based on pollution-tolerant taxa and uses a concept of watershed protection that emphasizes all aspects of water quality (Barbour et al., 1999). Its result corroborated what we found for the number of individuals and community composition, which is a gradation with the reference stream presenting the best environmental quality, the urban stream in an intermediate position and the agricultural stream with the worst environmental quality (Table 2).

The same pattern is verified by cluster analysis and by non-metric multidimensional scaling. In cluster analysis, we found a wide variation among the replicates of the same streams. However, anthropic changes are evident with the formation of three groups preferably separated by the type of disturbance assessed (Figure 3A). The NMDS displays the intermediate condition of the urban stream since the agricultural and the reference streams appear entirely separate, with the urban stream partially overlapping both (Figure 3B). In both analyses, it is possible to verify that cluster separation is not very noticeable and that streams still have overlapped communities. We already expected this effect since we evaluated streams subjected to low-intensity impacts in micro watersheds that have the same formation. This relationship is likely to be linear. That is, the more intense the disturbances, the longer the separation between the clusters. On the other hand, the absence of impacts would maintain replicas of different streams in the same group. Further studies would be required to confirm the linearity of disturbance intensity and the interference of stream formation in the clustering. However, if linearity is confirmed, and interference is negligible, aquatic insect community cluster analysis can be used to identify priority areas for environmental restoration (i.e., areas that are totally separate from their references) and also to measure the effectiveness of these restoration actions (i.e., how close the area has come to the reference).

As mentioned earlier, any study related to the subsidy-stress gradient should make it very clear about the scale and classification of ecosystem responses to environmental changes. In our case, we understand that specific changes in processes are less significant than the ecosystem’s integrated response to the disturbance (macro-scale) and that quantification of process yields are less significant than the understanding of the effect of that process on the ecosystem homeostasis (classification). For example, we consider the increase in community respiration leading to a process of eutrophication (Pereda et al., 2019) as part of a stress effect, and the burning of an area of vegetation leading to a regeneration process (Durigan & Ratter, 2016) as part of a subsidy effect.

Therefore, we conclude that the response of headwater streams to the mild impacts of both agricultural activity and urban settlement can be characterized as stress effects, although these effects have different characteristics. The stress effect in the agricultural stream is as hypothesized by the subsidy-stress gradient, a response consistent with the input of toxic pollutants into the ecosystem even at low intensity. However, what we observed in the urban stream was not an increase on ecosystem performance but an imbalance. Although it is not strong enough to bring down measures of richness and diversity, this imbalance affects the composition and density of the aquatic insect community, and therefore we can not consider it as a subsidy effect.

## Supporting information

All supplemental material

## Acknowledgements

We thank Leandro Lourenço Dumas and Ana Lúcia Oliveira for their contributions in the identification of organisms, Nicolas J. Locke for the opportunity to develop research at the Guapiaçu Ecological Reserve (REGUA) and the UFRJ Zoology and Ecology postgraduate programs, who funded our stay at REGUA.

