## Supplementary material for "Mild impacts do not subsidize aquatic insect communities in an Atlantic Forest stream": All supplemental material

### Supplementary Material 1 – Raw data and calculation codes

```
### Pacotes utilizados
suppressPackageStartupMessages(library("vegan"))
suppressPackageStartupMessages(library("SPECIES"))
suppressPackageStartupMessages(library("ineq"))
suppressPackageStartupMessages(library("dendextend"))
suppressPackageStartupMessages(library("ggplot2"))
suppressPackageStartupMessages(library("ggrepel"))
suppressPackageStartupMessages(library("rareNMtests"))

### Função de cálculo de alfa diversidade
alpha.div=function(a) {
 c=subset(a, apply(a,1,sum)>0) # Seleciona valores positivos da otu table
 d=apply(c,1,sum) # Calcula a abundância por espécie.
 e=as.vector(d) # Transforma o dado em um vetor.
 f=length(e) # Calcula a riqueza de otus na amostra.
 b=sum(e)
 g=sum(e==1) #singletons
 h=sum(e==2) #doubletons
 i=round(diversity(e),2) #Shannon
 j=round(fisher.alpha(e),2) #Alfa de Fisher
 k=round(i/log(length(e)),2) #Pielou
 l=round(ineq(e, "Gini"),2) #Gini
 m=table(e)
 n=as.numeric(as.vector(rownames(m)))
 o=as.numeric(as.vector(m))
 p=as.matrix(cbind(n,o))
 q=chao1984(p)
 r=q$Nhat #Chao1 para tabela de abundância
 #r.low=q$CI[1,1] #Chao1 lower bound
 #r.up=q$CI[1,2] #Chao1 upper bound
 s=jackknife(p, k=2)
 t=s$Nhat[1] #Jackknife de segunda ordem
 #t.low=s$CI[1,1] #Jackknife lower bound
 #t.up=s$CI[1,2] #Jackknife upper bound
 u=ChaoLee1992(p, t=round(sum(e)/length(e)), method="ACE")
 v=u$Nhat #Estimador ACE
 #v.low=u$CI[1,1] #ACE lower bound
 #v.up=u$CI[1,2] #ACE upper bound
 w=cbind("Indivíduos"=b, "Riqueza"=f, "Singles"=g, "Doubles"=h, "Chao1"=r, "Jack2"=t,
"ACE"=v,
"Shannon"=i, "F.Alfa"=j, "Pielou"=k, "Gini"=l)
 rownames(w) = deparse(substitute(a)) #Coloca o nome do objeto de entrada como nome da linha.
 return(w)
 }

### Dados brutos
ia.raw=matrix(c(0, 0, 0, 0, 0, 0, 0, 0, 0, 0, 0, 0, 0, 0, 3, 0, 0, 0, 2, 0, 0, 0, 0, 0, 0, 0, 0, 0, 0, 0, 5, 13,
59, 43, 1, 2, 15, 2, 4, 5, 2, 16, 1, 1, 1, 0, 0, 4, 42, 0, 0, 0, 0, 1, 0, 0, 1, 3, 0, 5, 0, 0, 0, 0, 0, 0, 0, 0, 0,
0, 0, 0, 1, 0, 0, 0, 0, 0, 0, 0, 0, 0, 1, 0, 0, 0, 0, 0, 0, 0, 0, 1, 0, 1, 0, 0, 0, 1, 0, 0, 0, 0, 2, 0, 0, 0, 0, 0,
0, 0, 0, 0, 0, 0, 0, 1, 0, 0, 0, 13, 1, 26, 320, 6, 20, 37, 15, 64, 24, 49, 67, 53, 15, 37, 0, 2, 0, 8, 0, 0, 0,
```

```
0, 0, 0, 0, 0, 1, 0, 1, 1, 0, 0, 0, 0, 0, 0, 0, 0, 0, 0, 0, 0, 0, 0, 0, 2, 0, 0, 1, 0, 0, 0, 0, 0, 0, 0, 0, 0, 0, 0, 0,
0, 0, 0, 0, 0, 0, 0, 0, 0, 2, 0, 0, 0, 0, 0, 1, 0, 0, 0, 0, 0, 0, 0, 0, 0, 0, 0, 0, 0, 0, 25, 58, 2, 11, 129, 0, 2,
31, 0, 18, 1, 1, 0, 0, 0, 0, 0, 0, 0, 0, 0, 0, 0, 0, 1, 0, 1, 0, 0, 0, 1, 0, 0, 0, 0, 0, 0, 0, 0, 0, 0, 0, 0, 0, 0, 0,
1, 0, 0, 0, 0, 0, 0, 0, 0, 0, 3, 0, 0, 0, 2, 0, 0, 0, 0, 0, 0, 0, 0, 0, 0, 0, 0, 2, 3, 2, 0, 7, 0, 0, 0, 1, 2, 0, 0, 0,
1, 0, 0, 0, 0, 0, 0, 0, 0, 1, 0, 0, 0, 0, 0, 0, 0, 0, 0, 5, 0, 5, 0, 0, 1, 0, 5, 0, 1, 4, 9, 4, 28, 9, 4, 0, 10, 0, 0,
1, 2, 2, 2, 17, 9, 13, 6, 0, 0, 0, 0, 0, 0, 0, 0, 0, 0, 0, 0, 1, 0, 0, 0, 0, 0, 0, 0, 0, 0, 1, 0, 0, 0, 0, 1, 1, 0, 0,
0, 0, 0, 0, 0, 0, 0, 0, 1, 0, 0, 0, 0, 0, 0, 0, 0, 0, 0, 0, 0, 0, 0, 0, 0, 0, 1, 0, 0, 0, 0, 0, 0, 0, 0, 0, 2, 5, 0, 0,
0, 0, 0, 0, 0, 1, 0, 0, 0, 0, 0, 0, 0, 0, 0, 0, 0, 0, 0, 1, 0, 0, 0, 0, 0, 0, 0, 0, 0, 0, 0, 0, 0, 0, 0, 0, 0, 0, 0, 0,
0, 0, 0, 0, 0, 1, 1, 0, 0, 0, 0, 0, 0, 0, 1, 0, 0, 0, 0, 0, 0, 0, 0, 0, 0, 1, 1, 1, 0, 0, 0, 0, 0, 0, 0, 0, 0, 0, 0, 0,
6, 21, 4, 1, 0, 0, 0, 0, 0, 0, 0, 2, 7, 5, 0, 0, 0, 0, 0, 0, 0, 0, 0, 0, 0, 1, 0, 0, 0, 3, 18, 15, 4, 3, 0, 0, 0, 1,
0, 0, 0, 0, 0, 0, 4, 5, 2, 0, 0, 0, 0, 0, 0, 0, 0, 0, 1, 0, 0, 0, 2, 2, 1, 6, 38, 1, 0, 6, 0, 17, 19, 4, 25, 194,
89, 3, 0, 106, 9, 2, 4, 20, 5, 1, 1, 0, 2, 0, 0, 0, 5, 5, 0, 0, 3, 0, 0, 2, 0, 30, 10, 9, 56, 0, 0, 1, 0, 0, 0, 0,
0, 0, 0, 0, 0, 0, 0, 0), 41, 15, byrow=TRUE)
rownames(ia.raw)=c("Blaberidae", "Driopidae", "Elmidae", "Elmidae_(A)", "Psephenidae",
"Staphylinidae", "Ceratopogonidae", "Ceratopogonidae_(P)", "Chironomidae",
"Chironomidae_(P)", "Culicidae", "Dixidae", "Empididae", "Ephydridae", "Simulidae",
"Simulidae_(P)", "Syrphidae", "Tabanidae", "Tipulidae", "Baetidae", "Caenidae", "Leptohyphidae",
"Leptophlebiidae", "Mesoveliidae", "Naucoridae", "Pleidae", "Lepidoptera", "Crambidae",
"Calopterigidae", "Coenagrionidae", "Libellulidae", "Megapodagrionidae", "Gripopterigidae",
"Perlidae", "Trichoptera_(P)", "Calamoceratidae", "Helicopsychidae", "Hydropsichidae",
"Leptoceridae", "Philopotamidae", "Polycntropodidae")
colnames(ia.raw)=paste0(c(rep("RF",5), rep("AG",5), rep("UR",5)), rep(c(1:5),3))
ia.tax=c("Blattodea;Blaberidae", "Coleoptera;Driopidae", "Coleoptera;Elmidae",
"Coleoptera;Elmidae_(A)", "Coleoptera;Psephenidae", "Coleoptera;Staphylinidae",
"Diptera;Ceratopogonidae", "Diptera;Ceratopogonidae_(P)", "Diptera;Chironomidae",
"Diptera;Chironomidae_(P)", "Diptera;Culicidae", "Diptera;Dixidae", "Diptera;Empididae",
"Diptera;Ephydridae", "Diptera;Simulidae", "Diptera;Simulidae_(P)", "Diptera;Syrphidae",
"Diptera;Tabanidae", "Diptera;Tipulidae", "Ephemeroptera;Baetidae", "Ephemeroptera;Caenidae",
"Ephemeroptera;Leptohyphidae", "Ephemeroptera;Leptophlebiidae", "Hemiptera;Mesoveliidae",
"Hemiptera;Naucoridae", "Hemiptera;Pleidae", "Lepidoptera", "Lepidoptera;Crambidae",
"Odonata;Calopterigidae", "Odonata;Coenagrionidae", "Odonata;Libellulidae",
"Odonata;Megapodagrionidae", "Plecoptera;Gripopterigidae", "Plecoptera;Perlidae",
"Trichoptera_(P)", "Trichoptera;Calamoceratidae", "Trichoptera;Helicopsychidae",
"Trichoptera;Hydropsichidae", "Trichoptera;Leptoceridae", "Trichoptera;Philopotamidae",
"Trichoptera;Polycntropodidae")
ia=cbind(ia.raw, as.data.frame(ia.tax))
colnames(ia)[16]="taxonomy"

### Composição da comunidade
ia.comp=subset(ia, apply(ia[,1:15], 1, max) > 2)
ia.comp=read.csv("/home/biocelio/Pubs/ia_comp.csv", header=TRUE)
x11(width=10, height=10)
#tiff(filename="/home/biocelio/Pubs/ia/ia_comp_color.tiff", width=10, height=10, units="in",
res=500)
ggplot(ia.comp, aes(x=taxa, y=number, color=local, fill=local)) +
 scale_color_manual(values=c("orange2", "green4", "blue")) +
 scale_fill_manual(values=c("orange2", "green4", "blue")) +
 theme_light() +
 theme(text=element_text(size=18), legend.position = "none") +
 ylim(0,330) +
 ylab("") +
 xlab("") +
```

```
 geom_boxplot(alpha=0.70) +
 coord_flip()
dev.off()

### Dados por estação amostral
ia.rf=ia[,1:5]
ia.ag=ia[,6:10]
ia.ur=ia[,11:15]
ia.rf.01=apply(ia.rf, 1, sum)
ia.ag.01=apply(ia.ag, 1, sum)
ia.ur.01=apply(ia.ur, 1, sum)
ia.01=cbind("Referência"=ia.rf.01, "Agrícola"=ia.ag.01, "Urbana"=ia.ur.01)

### Alfa diversidade
alpha=t(rbind(alpha.div(ia.rf,480), alpha.div(ia.ag,480), alpha.div(ia.ur,480)))
alpha[alpha==Inf]=0

### Gráfico de Rarefação
rar.rf=rarefaction.individual(ia.01[,1])
colnames(rar.rf)=c("individuals", "richness")
rar.ag=rarefaction.individual(ia.01[,2])
colnames(rar.ag)=c("individuals", "richness")
rar.ur=rarefaction.individual(ia.01[,3])
colnames(rar.ur)=c("individuals", "richness")
x11(width=5, height=5, family="Courier New")
#tiff(filename="/home/biocelio/Pubs/ia/rarefaction.tiff", width=5, height=5, family="Courier New",
units="in", res=500)
ggplot() + ylim(0,27) +  theme_light() +
 geom_point(aes(x=individuals, y=richness), data=rar.rf, size=0.1, colour="green4") +
 geom_line(aes(x=individuals, y=richness), data=rar.rf, size=1, colour="green4") +
 geom_point(aes(x=individuals, y=richness), data=rar.ag, size=0.1, colour="orange2") +
 geom_line(aes(x=individuals, y=richness), data=rar.ag, size=1, colour="orange2") +
 geom_point(aes(x=individuals, y=richness), data=rar.ur, size=0.1, colour="blue") +
 geom_line(aes(x=individuals, y=richness), data=rar.ur, size=1, colour="blue") +
 geom_vline(xintercept = 480, color = "red", size=0.35) +
 theme(axis.text=element_text(size=12), axis.title=element_text(size=14,face="bold"))
dev.off()

### Gráfico de Acumulação
spc.rf=specaccum(t(ia.rf), method="exact")
spc.rf.df=cbind("samples"=spc.rf$sites, "richness"=as.numeric(spc.rf$richness))
spc.rf.df=as.data.frame(spc.rf.df)
spc.ag=specaccum(t(ia.ag), method="exact")
spc.ag.df=cbind("samples"=spc.ag$sites, "richness"=as.numeric(spc.ag$richness))
spc.ag.df=as.data.frame(spc.ag.df)
spc.ur=specaccum(t(ia.ur), method="exact")
spc.ur.df=cbind("samples"=spc.ur$sites, "richness"=as.numeric(spc.ur$richness))
spc.ur.df=as.data.frame(spc.ur.df)
x11(width=5, height=5, family="Courier New")
#tiff(filename="/home/biocelio/Pubs/ia/accumulation.tiff", width=5, height=5, family="Courier
New", units="in", res=500)
ggplot() + ylim(0,27) +  theme_light() +
```

```
 geom_point(aes(x=samples, y=richness), data=spc.rf.df, size=1, colour="green4") +
 geom_line(aes(x=samples, y=richness), data=spc.rf.df, size=1, colour="green4") +
 geom_point(aes(x=samples, y=richness), data=spc.ag.df, size=1, colour="orange2") +
 geom_line(aes(x=samples, y=richness), data=spc.ag.df, size=1, colour="orange2") +
 geom_point(aes(x=samples, y=richness), data=spc.ur.df, size=1, colour="blue") +
 geom_line(aes(x=samples, y=richness), data=spc.ur.df, size=1, colour="blue") +
 theme(axis.text=element_text(size=12), axis.title=element_text(size=14,face="bold"))
dev.off()

x11(width=7, height=7, family="Courier New")
#tiff(filename="/home/biocelio/ia/rarefaction.tiff", width=7, height=7, family="Courier New",
units="in", res=500)
rarecurve(t(ia.01), sample=480, xlab="Number of Individuals", ylab="Number of Families",
label=FALSE, col=c("gray80", "gray60", "gray40"), bty="l", lwd=4)
legend(x=800, y=9, legend="Referência", col="gray80", lty=1, lwd=6, bty="n")
legend(x=800, y=7, legend="Agrícola", col="gray60", lty=1, lwd=6, bty="n")
legend(x=800, y=5, legend="Urbana", col="gray40", lty=1, lwd=6, bty="n")
dev.off()

### EPT & LPN
ia.ept=ia[c(20:23,33:41),]
ia.ept.rf=ia.ept[,1:5]
ia.ept.ag=ia.ept[,6:10]
ia.ept.ur=ia.ept[,11:15]
ia.ept.rf.01=apply(ia.ept.rf, 1, sum)
ia.ept.ag.01=apply(ia.ept.ag, 1, sum)
ia.ept.ur.01=apply(ia.ept.ur, 1, sum)
EPT.rf=round(100 * (sum(ia.ept.rf.01>0)/sum(ia.rf.01>0)),2)
EPT.ag=round(100 * (sum(ia.ept.ag.01>0)/sum(ia.ag.01>0)),2)
EPT.ur=round(100 * (sum(ia.ept.ur.01>0)/sum(ia.ur.01>0)),2)
EPT=c(EPT.rf, EPT.ag, EPT.ur)
LPN=c(4.29, 5.42, 4.68) # From LPN reports
eptlpn=rbind(EPT, LPN)
colnames(eptlpn)=c("Referência", "Agrícola", "Urbana")

# NMDS
ia.nmds=ia[,1:15]
ia.nmds.01=t(ia.nmds)
ia.nmds.02=scale(ia.nmds.01, center=FALSE, scale=apply(ia.nmds.01, 2, sum))
ia.nmds.ord=metaMDS(ia.nmds.02)
ia.splabel=c("Bla", "Dri", "Elm", "ElmA", "Pse", "Sta", "Cer", "CerP", "Chi", "ChiP", "Cul", "Dix",
"Emp", "Eph", "Sim", "SimP", "Syr", "Tab", "Tip", "Bae", "Cae", "Lhy", "Lph", "Mes", "Nau",
"Ple", "Lpt", "Cra", "Cpt", "Coe", "Lib", "Meg", "Gri", "Per", "TriP", "Cmo", "Hel", "Hyd", "Lto",
"Phi", "Pol")

ia.nmds.pts=cbind(ia.nmds.ord$points[,1], ia.nmds.ord$points[,2],
as.data.frame(substr(names(ia.nmds.ord$points[,2]), start = 1, stop = 2)))
colnames(ia.nmds.pts)=c("pt1", "pt2", "gr")
set.seed(1)
x11(width=6, height=5.8)
#tiff(filename="/home/biocelio/Pubs/ia/nmds.tiff", width=6, height=5.8, family="Courier New",
units="in", res=500)
```

```
ggplot() + theme_light() + xlab("NMDS1") + ylab("NMDS2") + #ylim(-0.4,0.64) + xlim(-0.5,0.6) +
 geom_point(aes(x=pt1, y=pt2, color=gr), data=ia.nmds.pts, alpha=1, size=2, shape=19) +
 scale_color_manual(values=c("orange2", "green4", "blue")) +
 scale_fill_manual(values=c("orange2", "green4", "blue")) +
 stat_ellipse(aes(x=pt1, y=pt2, color=gr, fill=gr), data=ia.nmds.pts, type="norm", geom="polygon",
level=0.67, alpha=0.25) +
 geom_point(aes(x=ia.nmds.ord$species[,1], y=ia.nmds.ord$species[,2]), alpha=1,
colour="gray50", size=1, shape=8)+
 geom_text(aes(x=pt1, y=pt2), data=ia.nmds.pts, label=rownames(ia.nmds.pts), colour="black",
hjust=c(-0.2,1.25,0.2,1.25,-0.2,-0.2,-0.2,-0.2,-0.2,1.25,-0.2,1.25,1.25,-0.3,1.2), vjust=c(-0.2,-
0.4,1.7,-0.2,-0.2,-0.2,-0.2,-0.2,-0.2,-0.2,-0.2,-0.2,-0.2,-0.2,1.4)) +
 geom_text_repel(aes(x=ia.nmds.ord$species[,1], y=ia.nmds.ord$species[,2]), label=ia.splabel,
colour="gray50", force=3, segment.size=0.15) +
 coord_fixed()
dev.off()

# Dendrogramas de Dissimilaridade
ia.den=ia[,1:15]
ia.den.01=t(ia.den)
ia.den.02=scale(ia.den.01, center=FALSE, scale=apply(ia.den.01, 2, sum))
ia.den.03=as.dendrogram(hclust(vegdist(ia.den.02)))
x11(width=4, height=5.8)
#tiff(filename="/home/biocelio/Pubs/ia/dendro.tiff", width=4, height=5.8, family="Courier New",
units="in", res=500)
par(mfrow=c(1,1), mar=c(4.5,0.5,0,2), cex=1)
ia.den.03 %>% set("branches_lwd", 3) %>% set("labels_cex", 1) %>% set("leaves_pch", ">") %>
% set("leaves_cex", 1) %>% set("leaves_col", "black") %>% plot(horiz=TRUE, xlab="Bray-Curtis
Dissimilarity", xlim=c(1.0,0.0))
dev.off()
```

**Supplementary Material 2 – Leaf Package Network reports**

Reference stream

### Calculated Values

Total Number of Individuals

241.00000000000003

Total Tolerance Value

1,033.4

Your Biotic Index

4.29

Your Percent EPT

47.1

### What Do These Numbers Mean?

**Biotic Index** is a comparison of the abundance of taxa and their tolerance to environmental stress. The index can indicate organic and nutrient pollution. The lower the biotic index, the better the water quality.

| Biotic index | Water quality | Degree of organic pollution |
|---|---|---|
| < 3.75 | Excellent | Organic pollution unlikely |
| 3.75–5.0 | Good | Some organic pollution |
| 5.1–6.5 | Fair | Substantial pollution likely |
| 6.6–10.0 | Poor | Severe pollution likely |

**Percent EPT** is short for the total number of Ephemeroptera (mayflies), Plecoptera (stoneflies), and Trichoptera (caddisflies). Many species within these three groups are sensitive to changes in water quality.

In general, the more EPT taxa, the better the water quality. There are exceptions: certain netspinning Trichoptera (*Hydropsychidae*) are more tolerant to pollution than other caddisflies and are therefore not counted as part of the total Percent EPT.

Results from the Leaf Pack Network® Biotic and Water Quality Calculator, https://leafpacknetwork.org/biotic-index/

Print your results

Previous

Next

### Calculated Values

Total Number of Individuals

96.80000000000001

Total Tolerance Value

452.7

Your Biotic Index

4.68

Your Percent EPT

39.3

---

### What Do These Numbers Mean?

**Biotic Index** is a comparison of the abundance of taxa and their tolerance to environmental stress. The index can indicate organic and nutrient pollution. The lower the biotic index, the better the water quality.

| Biotic index | Water quality | Degree of organic pollution |
|---|---|---|
| < 3.75 | Excellent | Organic pollution unlikely |
| 3.75–5.0 | Good | Some organic pollution |
| 5.1–6.5 | Fair | Substantial pollution likely |
| 6.6–10.0 | Poor | Severe pollution likely |

**Percent EPT** is short for the total number of Ephemeroptera (mayflies), Plecoptera (stoneflies), and Trichoptera (caddisflies). Many species within these three groups are sensitive to changes in water quality.

In general, the more EPT taxa, the better the water quality. There are exceptions: certain netspinning Trichoptera (*Hydropsychidae*) are more tolerant to pollution than other caddisflies and are therefore not counted as part of the total Percent EPT.

Results from the Leaf Pack Network® Biotic and Water Quality Calculator, https://leafpacknetwork.org/biotic-index/

Print your results

Previous
Next

Agricultural Stream

### Calculated Values

Total Number of Individuals

95.39999999999999

Total Tolerance Value

517.4

Your Biotic Index

5.42

Your Percent EPT

12.8

---

### What Do These Numbers Mean?

**Biotic Index** is a comparison of the abundance of taxa and their tolerance to environmental stress. The index can indicate organic and nutrient pollution. The lower the biotic index, the better the water quality.

| Biotic index | Water quality | Degree of organic pollution |
|---|---|---|
| < 3.75 | Excellent | Organic pollution unlikely |
| 3.75–5.0 | Good | Some organic pollution |
| 5.1–6.5 | Fair | Substantial pollution likely |
| 6.6–10.0 | Poor | Severe pollution likely |

**Percent EPT** is short for the total number of Ephemeroptera (mayflies), Plecoptera (stoneflies), and Trichoptera (caddisflies). Many species within these three groups are sensitive to changes in water quality.

In general, the more EPT taxa, the better the water quality. There are exceptions: certain netspinning Trichoptera (*Hydropsychidae*) are more tolerant to pollution than other caddisflies and are therefore not counted as part of the total Percent EPT.

Results from the Leaf Pack Network® Biotic and Water Quality Calculator, https://leafpacknetwork.org/biotic-index/

Print your results

Previous
Next
